# Associative emotional memory encoding: Insights from network stability analysis of an fMRI-driven bilinear dynamics

**DOI:** 10.64898/2026.01.27.701503

**Authors:** Weronika Dziarnowska, Melis Orhun, Yannan Zhu, Nils Kohn, Guillén Fernández, Matin Jafarian

## Abstract

The interplay between emotion and memory is a central topic in cognitive neuroscience, with open questions about the underlying neuronal mechanisms. This article studies dynamic interactions among the hippocampus, amygdala, and orbitofrontal cortex during an fMRI associative memory encoding task. Participants were clustered into three condition groups: Neutral–Neutral, Neutral–Emotional, or Emotional–Emotional, and viewed image pairs associated with their assigned condition. Using the dynamic causal modeling framework, we explore several dynamic models and show that a stochastic bilinear state-space model best describes the neuronal dynamics in all conditions. Furthermore, we use graph and control theory techniques to both validate and analyze the model. In particular, we analyze the network dynamics of each condition using tools from graph theory and stability theory and discuss the differences in the strength and direction of connectivity as well as the stability of each of these networks. We confirm the prior finding that memory is enhanced in emotional conditions, in particular in the Neutral-Emotional condition. In our work, this enhanced memory is associated with increased hippocampus–amygdala coupling and overall network connectivity. In addition, we show that in the Emotional–Emotional condition, the coupling of the hippocampus and amygdala, as well as the whole network connectivity increases when the first image’s valence is substantially less negative rated than the second image. This pattern mirrors the Neutral–Emotional condition, where the first image is neutral compared with the second one. Moreover, our model-based analyses suggest that the amygdala predominantly influences the other two regions in the Neutral–Emotional condition, whereas the OFC plays a dominant role in the other two conditions. Combined data-driven modeling, stability analyses, and graph-theory tools led to new insights and enhanced the mechanistic understanding of cortical dynamics of emotional associative memory. We discuss these insights, utilize these analytical tools to generalize our findings to some unmeasured conditions, and highlight the potential of these techniques to inform the design of future regulatory mechanisms.

## 1 Introduction

The intricate relationship between emotions and memory has long fascinated researchers due to its profound implications for understanding human cognition and behavior, central to cognitive neuroscience and psychiatric research [33]. Emotional experiences are often remembered more vividly than neutral ones, shaping how individuals perceive and interact with the world [5, 23]. Yet, the neural mechanisms that govern emotional associative memory remain partially understood.

The brain regions involved in memory-emotion interactions include the amygdala, the hippocampus, and specific prefrontal areas. The amygdala is mainly involved in assigning emotional value to sensory stimuli and in modulating memory consolidation and retrieval [31] while the hippocampus supports the formation and retrieval of declarative, associative memory [11]. The orbito-frontal cortex (OFC) within the PFC exerts top-down control by integrating emotion regulation, attention, working memory, and reward processing to support flexible, goal-directed behavior [30]. To reveal the mechanisms of brain networks in cognition, data collected by functional magnetic resonance imaging (fMRI) has often been used. The technique offers non-invasive high-resolution imaging capable of capturing all brain regions, their structures, connectivity, and activity. Although some interactive mechanisms have been delineated, a comprehensive understanding has not yet been achieved.

To identify the underlying neural dynamics, several data-driven approaches have been discussed in the literature, among which are generative models which are a set of differential equations or density dynamics mainly represented as state-space models [35]. These models allow one to study interactions dynamically, enable mechanistic understanding from observations, and provide a basis for brain simulation and stimulation. There are various generative models depending on the level of abstraction, biological details, the amount of prior knowledge necessary, and the size of the datasets [2, 21]. Detailed biophysical models require extensive prior knowledge and are very complex in how they capture biological processes [20]. As greater complexity hinders parameterization, analysis, and inference, data-driven methods that require fewer biological assumptions are useful in revealing cortical level mechanisms [6, 9]. These approaches use techniques, such as Dynamic Causal Modeling (DCM), to estimate both the parameters of partially known biophysical models and the remaining model structure.

Dynamic Causal Modeling, in particular, has been known as a framework for modeling effective connectivity, that is, the causal influence that neuronal systems exert on each other [16]. It models the time evolution of latent neuronal states and hemodynamics assuming priors, and directly estimates endogenous, modulatory and driving influences within the target network. DCM has been extensively used in studying the effective connectivity of brain networks in several studies [7,15,25,26] with some of them introducing stochasticity [24], excitatory and inhibitory populations [28] and a nonlinear formulation [38].

Prior work has utilized DCM to study emotional memory mainly focusing on the amygdala and hippocampus interactions. One study, using deterministic two-state bilinear DCM, reported that the strength of the connection from the amygdala to the hippocampus was rapidly and robustly increased during the encoding of emotional, both positive and negative, pictures compared to the neutral ones [13]. Another fMRI study used classical DCM to study the effective connectivity of amygdala-hippocampus-dorsolateral prefrontal cortex in a memory and emotion task [18]. Their analysis showed that the suppression of distressing memories requires PFC regions to inhibit both amygdala and hippocampus activities. Using classical DCM, a further study on emotional associative memory retrieval demonstrated that the OFC modulates amygdala-hippocampus interactions during the recall of emotional context [36].

These studies demonstrate the utility of DCM in revealing effective connectivity in brain regions involved in emotional memory. To date, to the best of our knowledge, the obtained models have been mainly used for inferring effective connectivity and not for qualitative analysis of the network behavior. In fact, differential equations representing the emotional memory dynamics are capable of providing insights on the whole network performance and can be used for prediction and regulation. In this work, we aim at addressing this gap by modeling and analyzing the emotional memory encoding network in a task where the order and valence of negative emotions matter.

The recent study in [39] has performed fMRI data analysis from an experiment involving the memorization of neutral and emotionally charged image pairs and explored how emotional stimuli influence memory integration. Participants were clustered to three condition groups and viewed image pairs that were either Neutral–Neutral, or Neutral–Emotional, or Emotional–Emotional. The study has used task-dependent functional connectivity that allows measuring correlated activities among brain regions. Their findings suggested that emotional information facilitated memory integration with related neutral information but disrupted the integration with other emotional information.

In this work, we develop a dynamic model of the emotional associative memory encoding task reported in [39]. Our aims are to: (1) use Dynamic Causal Modeling (DCM) to derive a set of differential equations, i.e. dynamic model, that reproduce the data and captures the underling neuronal dynamics of emotional associative memory; (2) validate the dynamical properties of the model through stability and controllability analyses; (3) infer effective connectivity among the amygdala, hippocampus, and orbitofrontal cortex (OFC); (4) apply graph-theoretic measures to assess network connectivity and neuronal synchrony; and (5) evaluate how emotional valence, as the model input, affects the model’s dynamic properties, particularly stability. We expect the model to generalize to some unmeasured conditions. We discuss the results, limitations, and directions for future research.

## 2 Materials and Methods

This section details the experimental paradigm, data acquisition and processing procedures, modeling framework and the model space employed to obtain the associative memory encoding dynamics.

### 2.1 Experimental Paradigm

The data used in this research have been derived from the associative encoding phase of a functional magnetic resonance imaging (fMRI) experiment designed to investigate how emotional information influences associative memory [39]. During this phase, participants were shown 48 different “ABC” image triplets, where a single spatial location cue (A) was paired first with one image (B), and then the same location was paired with a second image (C), forming AB and AC pairs. The images B and C carried emotional valence; either neutral or negative. Participants were instructed to vividly imagine the relationship between the location and the associated image to facilitate memory formation.

A total of 70 healthy young adults completed the experiment. They were randomly assigned to one of three condition groups, based on the emotional valence of the image stimuli they were exposed to:

- **Neutral–Neutral (NN) group with 25 participants:** both associated images (B and C) in a triplet were neutral,
- **Neutral–Emotional (NE) group with 21 participants:** the first image (B) was neutral and the second image (C) was emotional (negative),
- **Emotional–Emotional (EE) group with 24 participants:** both images (B and C) were emotional (negative).

The 48 ABC triplets were split into four sets of 12 triplets. During the study, each participant completed four runs of the experiment, with a different set of 12 used per run. In each run, the AB and AC pairings were displayed in a blocked and repeated manner: 12 AB pairs were shown in consecutive encoding trials (explained below), then 12 AC pairs, followed by a repetition of the same AB and AC pairs, concluding a run. Throughout this process, the participants were actively monitored using fMRI and the BOLD responses were recorded. As shown in Figure 1, each encoding trial (AB or AC) consisted of:

**Figure 1.**
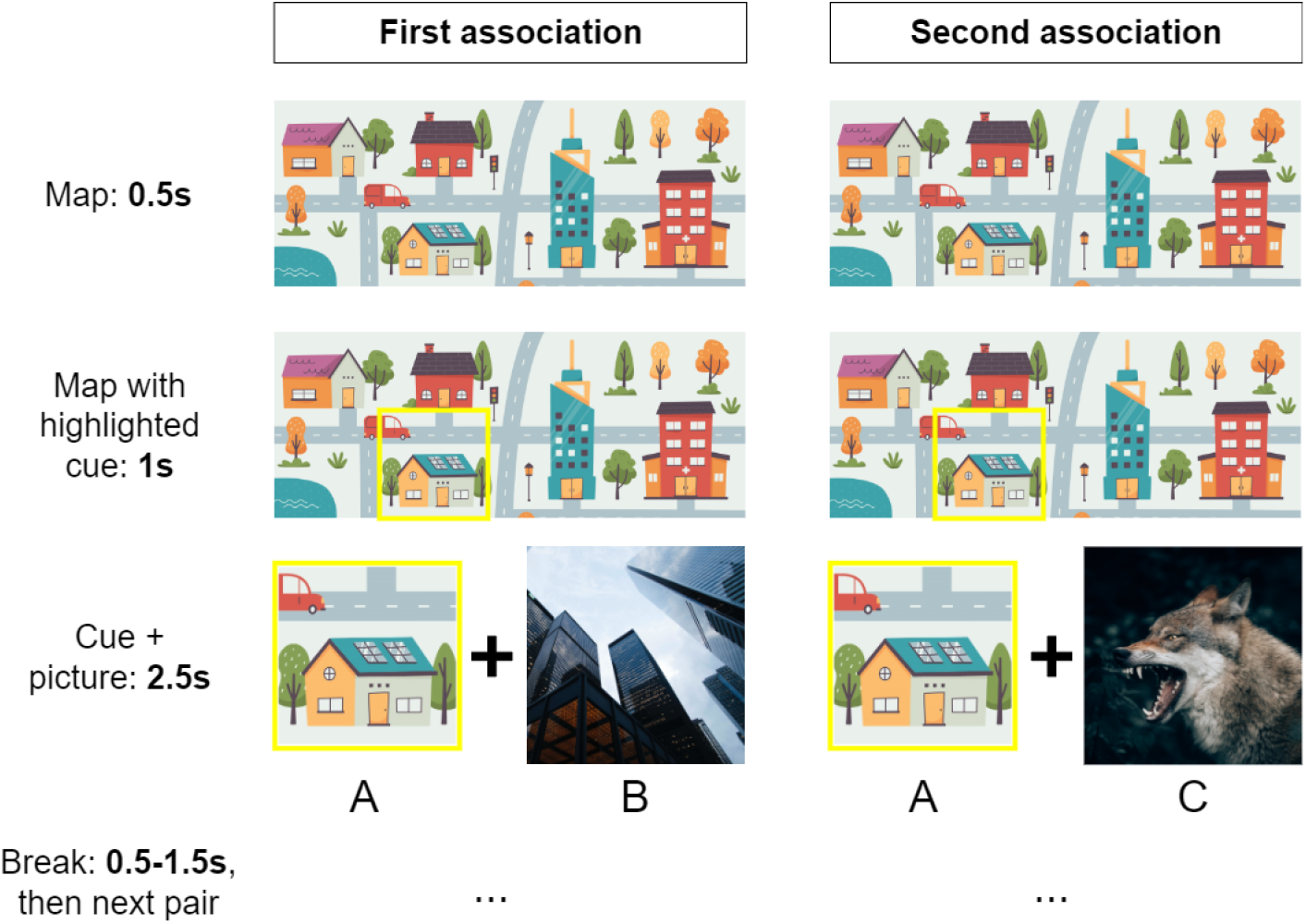
Associative encoding trial of the experiment. Adopted from [39]

1. A brief display of a cartoon map (0.5 s),
2. A highlighted location on that map (1.0 s),
3. A simultaneous display of the location cue (A) and the associated item (B or C) (2.5 s).

Trials were separated by a jittered inter-trial interval ranging from 0.5 to 1.5 seconds in which a fixation cross was displayed. As a result, each trial lasted approximately 5 seconds and each run around 4 minutes and 30 seconds. The experiment procedure also included recall and behavioral analysis phases. However, this study focuses exclusively on the encoding phase, as it provides the most direct insight into the neural mechanisms of emotional associative memory formation.

#### 2.1.1 Data acquisition

MRI data were acquired using a 3.0 T Siemens Skyra (Siemens Medical, Erlangen, Germany) with a 32-channel head coil system at the Donders Institute, Centre for Cognitive Neuroimaging in Nijmegen, the Netherlands. Functional images were collected using a multi-band echo-planar imaging (mb-EPI) sequence (slices, 66; multi-slice mode, interleaved; slice thickness, 2 mm; TR, 1000 ms; TE, 35.2 ms; flip angle, 60°; multiband accelerate factor, 6; voxel size, 2 × 2 × 2 mm; FOV, 213 × 213 mm). To correct for spatial distortions, fieldmap images were acquired (slices, 66; multi-slice mode, interleaved; slice thickness, 2 mm; TR, 500 ms; TE1, 2.80 ms; TE2, 5.26 ms; flip angle, 60°; voxel size, 2 × 2 × 2 mm; FOV, 213 × 213 mm). Structural images were acquired using a three-dimensional sagittal T1-weighted magnetization-prepared rapid gradient echo (MPRAGE) sequence (slices, 192; slice thickness, 1 mm; TR, 2300 ms; TE, 3.03 ms; flip angle, 8°; voxel size, 1 × 1 × 1 mm; FOV, 256 × 256 mm).

### 2.2 DCM data processing

Here we explain preparing data to be used in DCM. For preliminary data preprocessing we refer to [39]. The fMRI dataset consists of voxels (units on a 3D grid) in the brain regions and a corresponding time series for each voxel. The goal of data processing is to identify the task-relevant voxels within specific brain regions (Volumes of Interest, VOIs), and then average their time series to obtain a single representative signal per VOI. The VOIs considered are the amygdala (Amy), hippocampus (Hip) and orbitofrontal cortex (OFC), restricted to the left hemisphere for simplicity. These anatomical regions are identified using the Automated Anatomical Labeling (AAL) atlas via the WFU PickAtlas toolbox in MATLAB, which generates binary brain masks.

Task-relevant voxels are identified by fitting a Generalized Linear Model (GLM) to the BOLD signal at each voxel. A voxel is considered significant if its activity correlates with the experimental design, here, showing significant responses to both inputs. The GLM is defined as [14]:

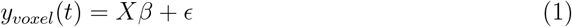

where *y*_*voxel*_(*t*) is the voxel’s observed BOLD signal over time, *X* is the design matrix, i.e., input signal convolved with a Hemodynamic Response Function(HRF), *β* are the parameter estimates, *ϵ* is the residual (unexplained) noise. The reduction of voxel data into single time series per VOI proceeds as follows:

1. **Define the design matrix** *X*: the external inputs are convolved with the canonical HRF.
2. **Apply anatomical brain masks:** Use WFU PickAtlas to restrict analysis to the left Amy, Hip, and OFC.
3. **Fit the GLM at each voxel to estimate** *β***s:** If the voxel’s time-series matches the predicted pattern, *β* will be significantly non-zero and the residual error will have low variance.
4. **Compute Tand F-contrasts to check significance of the voxels:** the significance of *β* is tested using:
  - T contrast tests whether a specific condition (e.g., each input) significantly activates the voxel.
  - F contrast tests whether a set of conditions has a significant joint effect.
5. **Extract a 1D time series per VOI:** average the time series of all significant voxels within each VOI.

We implemented the algorithm using the Statistical Parametric Mapping (SPM12) software package running in MATLAB [14]. At the end of pre-processing, participants with no significant voxels in at least one VOI were excluded leaving a final sample of 65 participants.

### 2.3 Modeling Framework

As motivated in the introduction, we chose a DCM framework for developing a state-space model that captures the dynamics of a three-region network comprising the Amygdala (Amy), Hippocampus (Hip), and Orbitofrontal Cortex (OFC). To this aim, we need to choose a model structure, e.g. bilinear, nonlinear, etc, external input signals, and the assumed connection across the network nodes, i.e., brain regions, as well as the manner that the exogenous inputs affect the connections or region dynamics. DCM estimates effective connectivity among brain regions using variational Bayesian inference under free-energy principle [16], briefly, a unified theory of how the brain combines prior knowledge with stimuli from the environment to learn and adapt.

The models’ outputs are the averaged time series per VOI obtained after the processing steps explained in Section 2.2. The inputs to our models are the deterministic signals defined as a step function takes the value 1 during external stimulation and 0 otherwise. As described in Section 2.1, each encoding trial has three stages: map, location cue, and simultaneous display of the item pair AB or AC. Because associative encoding is expected to occur only during co-presentation of AB or AC, the inputs are set to 1 only for this 2.5*s* and to 0 during the map, cue, and the jittered inter-trial interval. To distinguish first (AB) and second (AC) associations within each triplet, two separate input signals are defined: *u*_1_ for AB and *u*_2_ for AC. These functions are illustrated in Figure 2.

**Figure 2.**
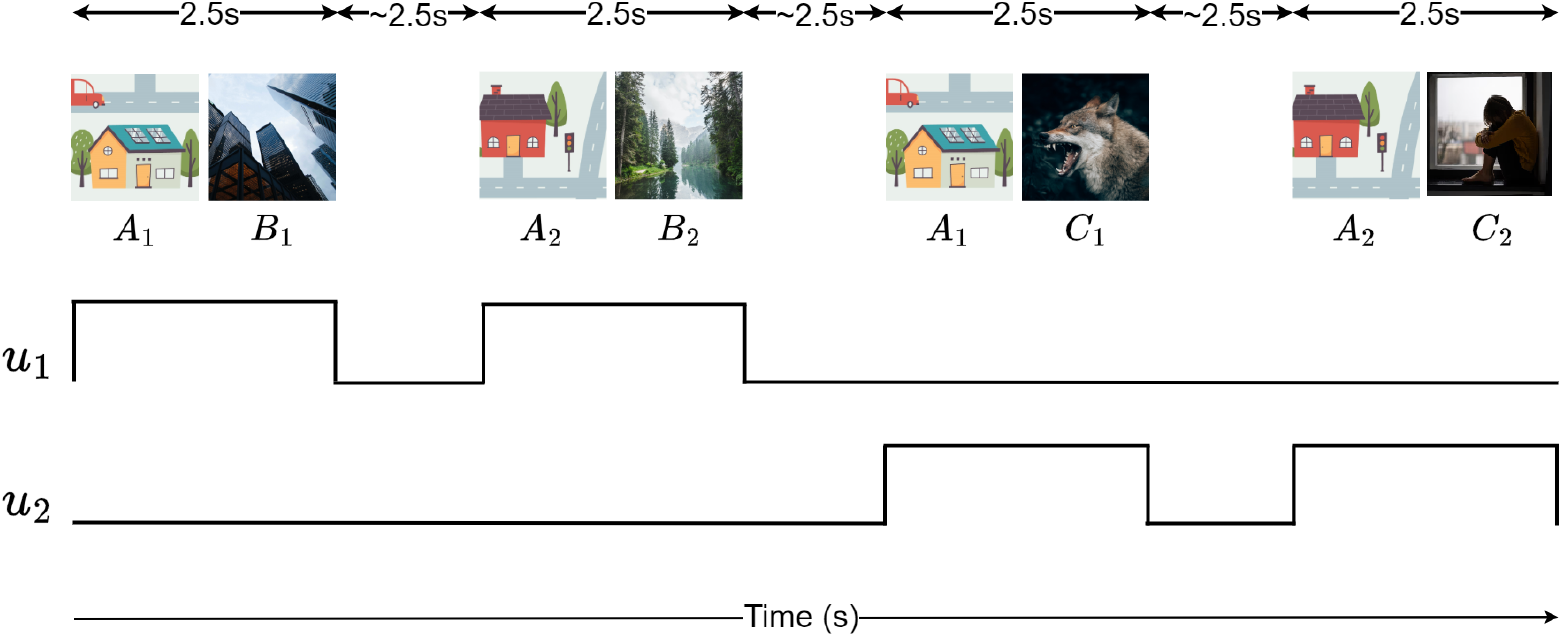
Example of input signals *u*_1_ and *u*_2_ used for fitting DCM in a dummy case with only two triplets in condition NE.

Models are identified using SPM12 on MATLAB. The algorithm expects the user to define which model connections are assumed. These connections are usually defined by using biological assumptions and results. If a connection is assumed, its value will be updated during model fitting, otherwise it will remain 0. In what follows, we provide a review on model structures, classic and stochastic DCM. An overview on nonlinear and two-state DCM, used in model comparison, is provided in Appendix.

#### 2.3.1 Classical (Bilinear) DCM

DCM modeling is composed of two dynamics: neuronal and hemodynamic states. For neuronal dynamics both model structure, inputs and parameters need to be identified, whereas the identification of hemodynamic only requires parameter identification. The neuronal dynamics are expressed by a nonlinear function that can be approximated by the Taylor Series expansion. Only the first-order derivatives are included in the classical DCM. The neuronal dynamics representing the Classical (Bilinear) DCM are:

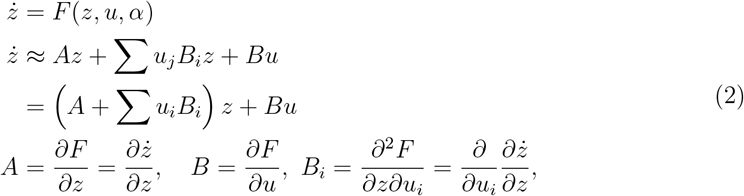

with *z* as states, *u* as inputs and *α* as the parameters of the model. Matrix *A* represents anatomical connections between the brain regions, *B*_*i*_ the change in coupling caused by the j-th input and *B* the direct influence of inputs on neuronal dynamics. Figure 3 shows the structure of the model.

**Figure 3.**
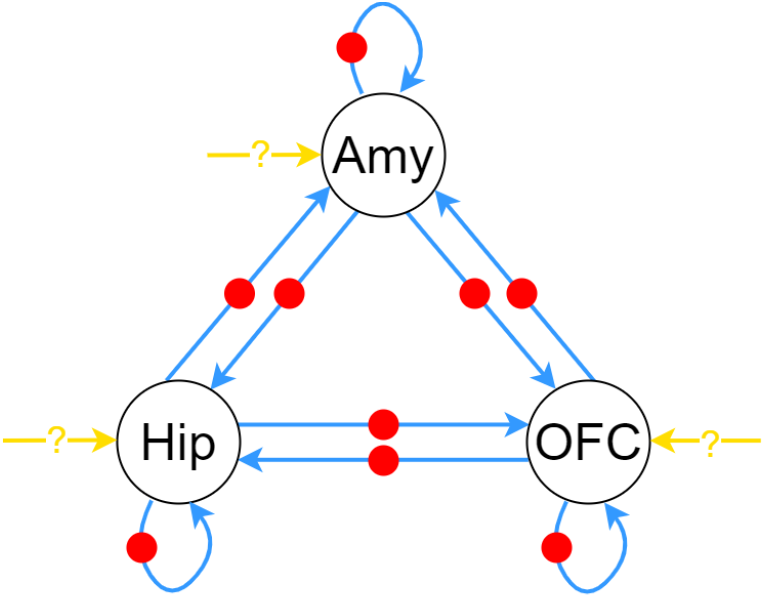
Model space for classical DCM. Blue arrows show couplings between nodes (*A*); dots show input-dependent modulation of couplings (*B*_*i*_); yellow arrow with question mark indicates direct driving effects of the external input on node states (*B*).

In addition to these dynamics, the DCM framework uses the Balloon model [3, 4, 27] for the hemodynamic which includes vasodilator signals, inflow, blood volume and normalized deoxyhemoglobin content. The dynamics explains how the activity of neural regions influence hemodynamic responses. When combined with the neuronal dynamics, the full model of the system is obtained. The final model then contains *x* as the states of both models, *u* the inputs and *θ* all of the parameters to be estimated for both models:

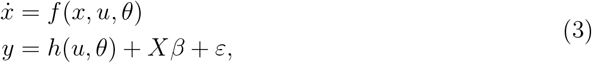

where *y* is the output, *h*(*u, θ*) is the estimated BOLD response, *X* captures the confounding effects, usually defined as a low-order discrete cosine that models low-frequency response drifts, with unknown coefficient *β* and *ε* is the error [16].

When using this formulation, noise and parameter priors are assumed Gaussian. The assumption is adopted to ease computation rather than to reflect biology [25]. With this model definition in mind, we choose the model structure of classical DCM as:

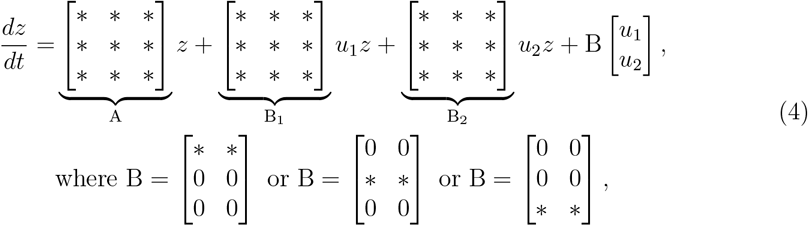

Where *denotes the parameter which needs to be identified. Here, all nodes are assumed to be connected with *A* matrix having all ones. This assumption is supported by previous studies on the connectivity of these brain regions [13, 18, 32, 36, 40]. Most of these studies found a bidirectional connection between the nodes which matches our assumptions for this matrix.

For the *B*_*i*_ matrices, we allow both inputs (*u*_1_, *u*_2_) to modulate all couplings, so differences between emotional and neutral inputs are expressed in the degree to which they modulate each connection. A similar assumption for *B*_*i*_ matrices have been imposed in another study on emotional associative memory [40].

In the case of matrix *B*, it is unclear which brain regions receive a direct input. However, related studies [13, 18, 32, 36, 40] found that the input acts directly on only one node which then influences the other nodes. Therefore, three different matrices are proposed, with inputs given solely to either Amy, Hip or OFC.

#### 2.3.2 Stochastic DCM

Deterministic variations of DCM omit the random firing of neurons and variations in transmissions between neurons due to the stochasticity at the cellular and molecular level [10, 12, 19, 37]. To account for these, stochastic DCM includes an additive term to model neuronal noise [8, 17, 24]. The resulting neuronal model, with *ϖ* as the noise, becomes:

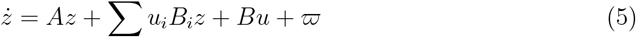

The final model is then obtained by including the same hemodynamic model as the other DCM formulations. In the end, the stochastic DCM includes two noise terms: the measurement noise *ε* which is common for all types of DCM utilized in this paper, and the neuronal noise *ϖ* introduced in the stochastic DCM. The node connections of Section 2.3.1 are kept for the stochastic DCM as well, yielding three models.

### 2.4 Model Fitting

In this research, the aim is to identify a single best-fitting DCM for each condition (NN, NE, EE) that explains the data across participants. To this end, we first fitted every model of the model space to each participant’s data for each of the 65 participants. By using four different model structures of DCM, we examined 18 models to be evaluated: 3 classical DCM, 9 nonlinear DCM, 3 two-state DCM and 3 stochastic DCM. The timing parameters for fitting were chosen to correspond to the experimental conditions. The echo time was to 40*ms* matching the property of the fMRI machine used [39]. In addition, slice-timing correction was applied to compensate for the delays within the Temporal Resolution (TR) of 1 sec. The time series were realigned to a reference time at the TR midpoint, following SPM12 recommendations. To fit the models, we used the data corresponding to first of four experimental runs per participants in [39]. Moreover, the first 10 initial values from each run has been discarded due to the scanner setup.

DCM models were estimated using variational Bayesian inference, which depends highly on prior specification. For all models, we adopted the default SPM12 priors without modification. After obtaining the subject-level models, we performed Bayesian Model Selection (BMS) separately for each condition (NN, NE, EE). Model comparison used log model evidence. We adopted fixed-effects BMS, which assumes a single best model for all participants corresponding to one condition given the identical task and conditions. Under this assumption, the winning model is common across participants, only the parameter estimates differ. Finally, we performed Bayesian Parameter Averaging (BPA) to obtain group-level parameter estimates for the winning model structure. This procedure was applied to each DCM type, classical, nonlinear, two-state, or stochastic, within each group yielding four optimal models for each of NN, NE and EE conditions.

### 2.5 Analytical tools

We now recall some of the tools from control theory [1, 22] and graph theory [29] applied in our analysis.

#### Control theory tools

- System 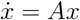 is asymptotically stable if and only if all the real part of all eigenvalues of matrix *A* are non-zero and negative.
- System 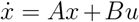 is controllable, meaning that the vector *x* can reach any desired vector in its space using control signal *u*, if and only if the controllability matrix *C* = {*B, AB*,…, *A*^*n−*1^*B*} is invertible.
- If the linearization of 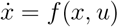 is controllable, then the nonlinear system is (locally) controllable.
- System 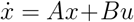 is Input-to-State Stable (ISS), i.e., bounded input gives bounded state *x*, if 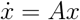 is asymptotically stable, and the input *u* is bounded.

#### Graph theory tools

The brain as a network represented by a graph is composed of brain regions, i.e., graph nodes, together with inter-region couplings, i.e., graph edges. Each node of the graph may have a connection to itself, i.e., self loop. The two nodes may be connected by one or two edges, directing to one another, whose weights can be different, i.e., weighted graph.

- The adjacency matrix of a directed weighted (digraph) graph captures the weights of all edges, as off-diagonal elements, as well as the self-loops, as diagonal elements.
- The graph Laplacian matrix of a digraph is *L* = *D*_in_ ^*−*^ *A*, where *A* is the adjacency matrix, and *D*_*in*_ is the diagonal in-degree matrix whose entries equal to the sum of weights of edges entering each node.
- Matrix *L* is positive semi-definite. The first non-zero eigenvalue of the graph Laplacian, the algebraic connectivity *λ*_2_, characterizes the network’s coordination. Larger *λ*_2_ implies stronger coupling and faster consensus, here synchrony in neuronal activities.

## 3 Results

This section first presents the result of model comparisons among four model structures. We show that the stochastic DCM performed best. Thereafter, we discuss the choices of external inputs, i.e. which of the three brain regions receives the external input per condition, and provide a comparison between the model output and the measured data. Next, we explore the system properties, stability and controllability, of the obtained models in order to validate them from a dynamical systems’ perspective.

### 3.1 Model Selection

We evaluated 18 mathematical models spanning classical, nonlinear, two-state and stochastic DCM formulations to explain effective connectivity within the Amy-Hip-OFC network during emotional associative encoding. To assess which of the models match the network best, we first performed Bayesian Model Selection (BMS) to determine the optimal model structure. Then, the parameters of this optimal model that explain all data belonging to a condition (NN, NE, EE) was obtained by Bayesian Parameter Averaging (BPA).

To quantify goodness of the fitting, we compute the coefficient of determination (*R*^2^) between the DCM-predicted time series and group-average measured data. The measure is used to compute fraction of variance in the measured time series that can be explained by the model, i.e.,

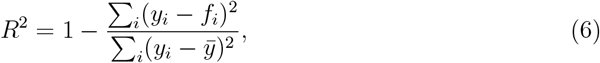

where *y*_*i*_ is the measured signal at time *i, f*_*i*_ is the model prediction and 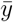 is the mean of the measured signal. Positive (*R*^2^) values closer to 1 indicate better fit, whereas negative values arise when the model performs worse than a mean-only (flat) fit. The results of this metric are displayed on Table 1. For classical, nonlinear and two-state models the (*R*^2^) values are negative or close to zero. In contrast, the stochastic DCM yields mostly positive (*R*^2^) with larger magnitudes, except for the OFC in the NE and EE conditions. These results, together with the qualitative observations from the figures, indicate that the stochastic DCM provides the best overall account of the data.

**Table 1.** The R-squared between the response of the optimal average models and the average measured data, calculated per condition, per brain region, and per DCM variation.

| | | $R^2_{\text{Amy}}$ | $R^2_{\text{Hip}}$ | $R^2_{\text{OFC}}$ |
| --- | --- | --- | --- | --- |
| **Classical DCM** | Condition NN | -6.7550 | -7.8580 | -3.1585 |
|  | Condition NE | -2.9134 | -2.0622 | -1.5884 |
|  | Condition EE | -3.3053 | -3.2494 | -0.0100 |
| **Nonlinear DCM** | Condition NN | -0.0411 | -0.8667 | -0.2948 |
|  | Condition NE | -0.0422 | -0.0120 | -0.0272 |
|  | Condition EE | 0.0075 | 0.0154 | 0.0005 |
| **Two-State DCM** | Condition NN | -0.0411 | -0.8667 | -0.2948 |
|  | Condition NE | 0.0241 | 0.0065 | 0.0088 |
|  | Condition EE | -1.9679 | -2.1009 | -0.0840 |
| **Stochastic DCM** | Condition NN | 0.6493 | 0.7626 | 0.3101 |
|  | Condition NE | 0.3889 | 0.2011 | -0.0344 |
|  | Condition EE | 0.4802 | 0.5599 | -0.6269 |

### 3.2 Stochastic DCM model: Structure and Accuracy

The stochastic DCM provided the best overall fit to the data, so we focus on this variant and examine its model structure, estimated parameters, state evolution properties and network connectivity.

#### Model structure: Bayesian Model Selection

As outlined in Section 2.3.1, we evaluate three stochastic models that share the same structure for *A* and *B*_*i*_ matrices of neuronal state equations but differ in their *B* matrices to identify which of the three nodes, Amy, Hip, and OFC, the external input stimulates. Bayesian Model Selection (BMS) compares candidate models using the log model evidence. This value combines how well a model fits the data with how simple it is, by penalizing deviations from priors. The log evidence values for the stochastic DCM models are reported in Table 2.

**Table 2.**
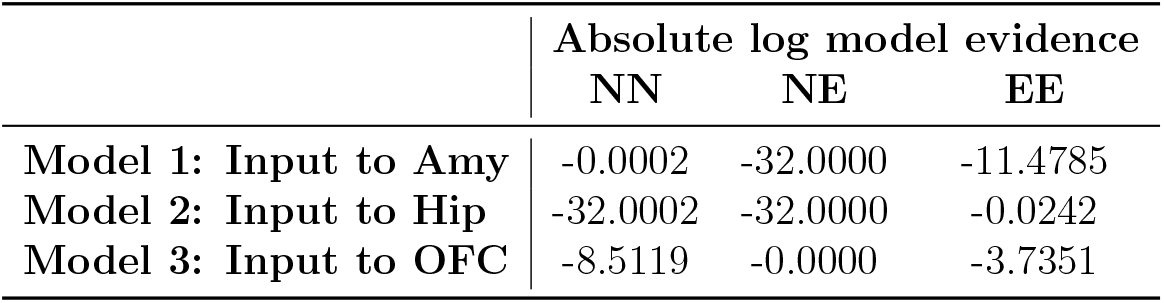
Absolute log model evidence values obtained during BMS.

It is easier to evaluate these values on a scale relative to the lowest value. The relative log evidence values are obtained by:

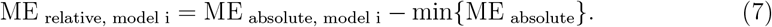

A relative log evidence of 3 or more is often considered as a strong indication for that model to be the optimal one. For each of the conditions, one of the models presents a relative log evidence of at least 3 compared to the other two models for that condition. These models are:

- Condition NN: Model 1 - input to Amy
- Condition NE: Model 3 - input to OFC
- Condition EE: Model 2 - input to Hip

Looking across the three conditions, the evidence is strongest in NE. In contrast, for EE, the comparison is least decisive since Models 2 and 3 separated by only a log evidence difference of just over 3. Notably, across all conditions, the OFC-driven model retains reasonable log evidence even when it is not the dominant one. This suggests a consistent role for OFC input in the encoding.

#### Model accuracy

Figure 4 shows the output of the group DCM model versus the data. In DCM for fMRI, neuronal states typically start at their prior mean, commonly zero, and hemodynamic states at the steady states. Initial states are not usually estimated as free parameters. For the stochastic DCM, although the response also starts at zero, it quickly shifts to the level set by the data and then tracks it without a sustained offset.

**Figure 4.**
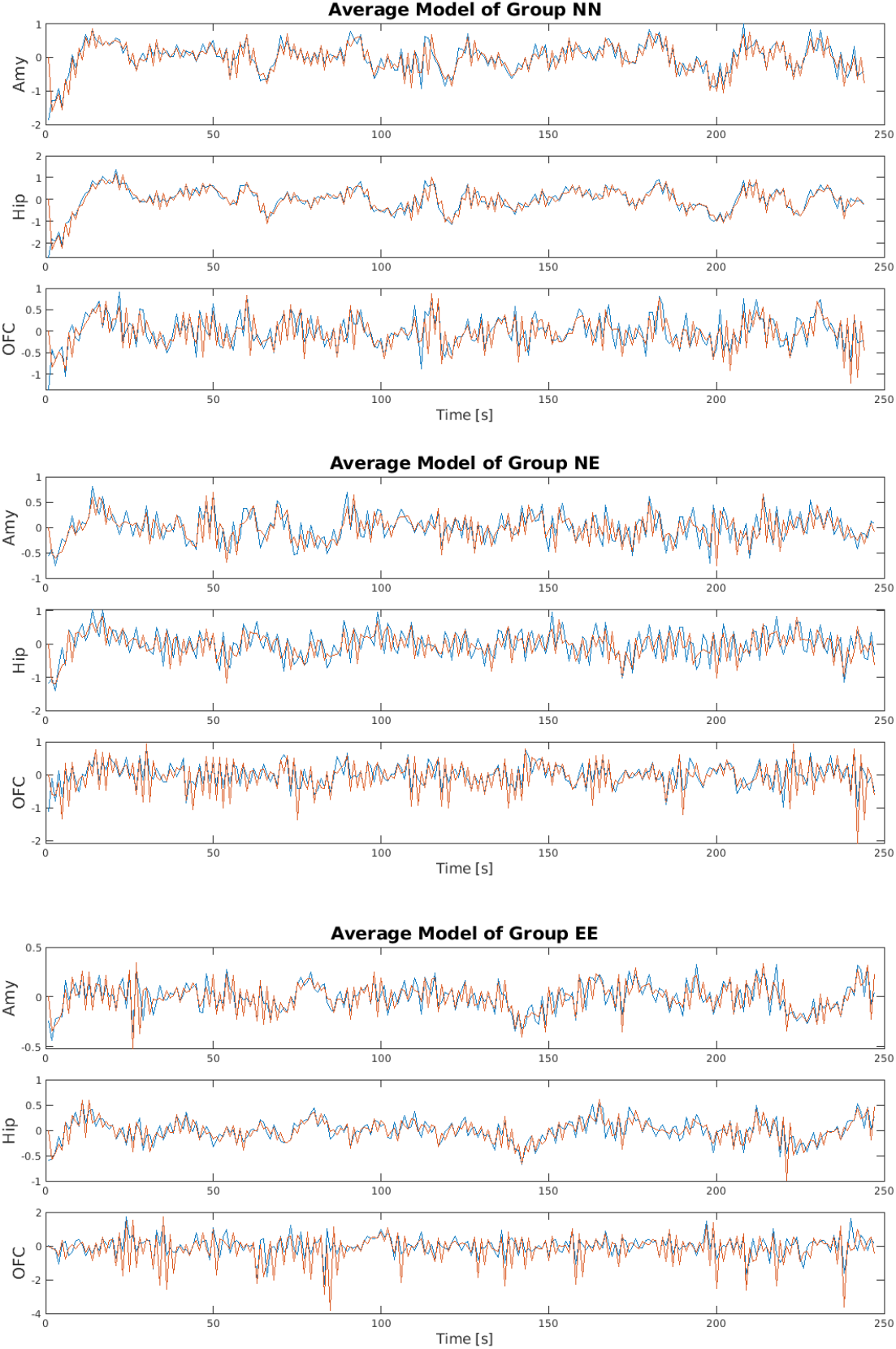
Stochastic DCM results for Amy, Hip and OFC: blue line is the response of the average optimal model obtained for each condition; red line is the average signal for each condition.

To assess node-wise fit, we examined coefficients of determination (*R*^2^) for all candidate models rather than only the BMS winners. Table 4 reports these values separately for each node under each input mapping for each condition, NN, NE, and EE. Higher *R*^2^ values indicate better fit to the data. Comparing the models selected by BMS, *R*^2^ is highest in the NN condition. This can be explained by emotional images introducing more complex, less predictable neural dynamics that are harder to capture.

**Table 3.** Number of significant voxels in the VOIs used to estimate group-optimal average models during BPA for all conditions.

|  | Number of significant voxels in BPA |  |  |
| --- | --- | --- | --- |
|  | Condition NN | Condition NE | Condition EE |
| Amy | 177 | 729 | 217 |
| Hip | 347 | 212 | 421 |
| OFC | 509 | 69 | 7 |

**Table 4.**
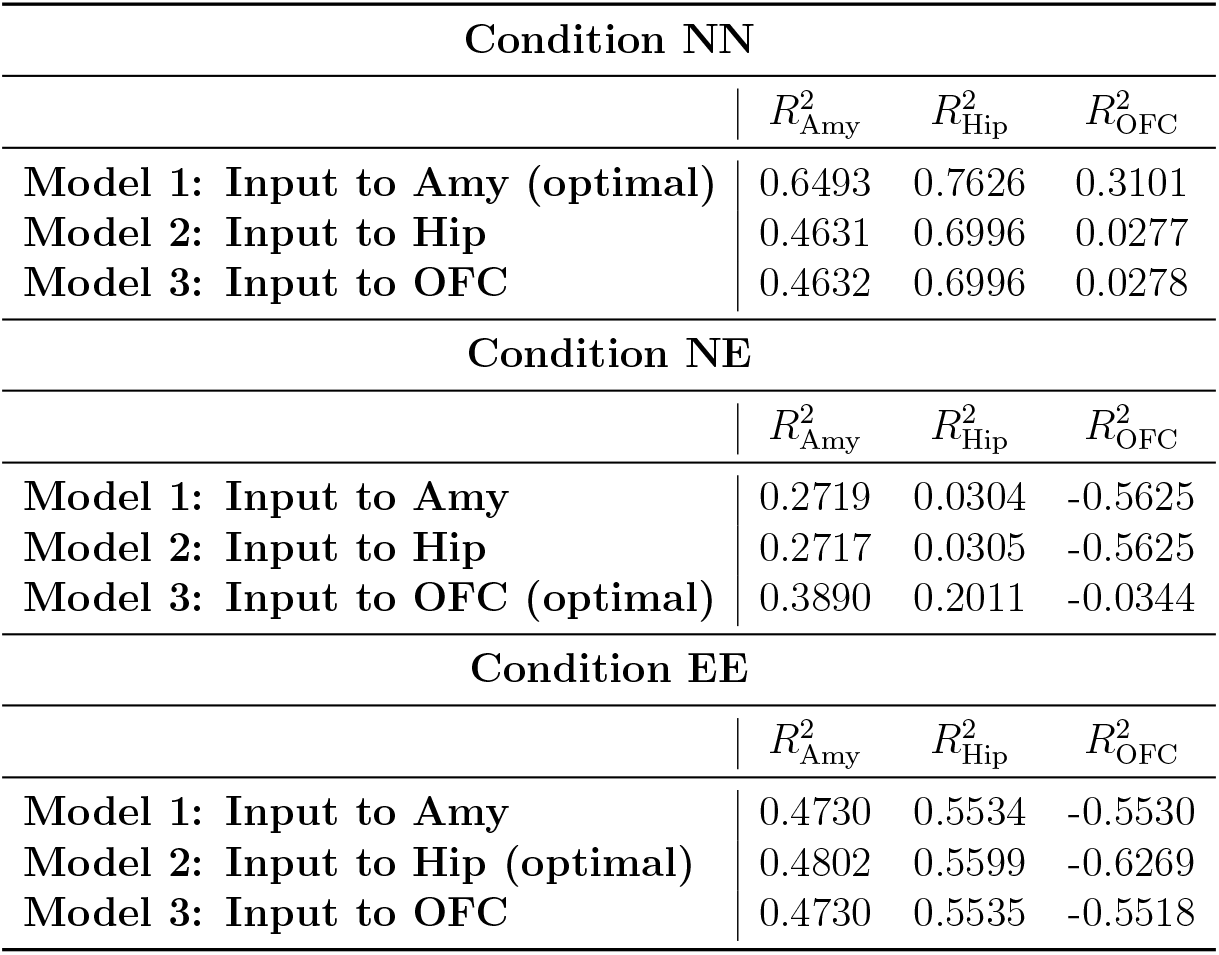
R-squared values in each node (Amy, Hip, OFC) for all average Stochastic DCM models.

| Condition NN |  |  |  |
| --- | --- | --- | --- |
| | $R^2_{\text{Amy}}$ | $R^2_{\text{Hip}}$ | $R^2_{\text{OFC}}$ |
| Model 1: Input to Amy (optimal) | 0.6493 | 0.7626 | 0.3101 |
| Model 2: Input to Hip | 0.4631 | 0.6996 | 0.0277 |
| Model 3: Input to OFC | 0.4632 | 0.6996 | 0.0278 |
| Condition NE |  |  |  |
| | $R^2_{\text{Amy}}$ | $R^2_{\text{Hip}}$ | $R^2_{\text{OFC}}$ |
| Model 1: Input to Amy | 0.2719 | 0.0304 | -0.5625 |
| Model 2: Input to Hip | 0.2717 | 0.0305 | -0.5625 |
| Model 3: Input to OFC (optimal) | 0.3890 | 0.2011 | -0.0344 |
| Condition EE |  |  |  |
| | $R^2_{\text{Amy}}$ | $R^2_{\text{Hip}}$ | $R^2_{\text{OFC}}$ |
| Model 1: Input to Amy | 0.4730 | 0.5534 | -0.5530 |
| Model 2: Input to Hip (optimal) | 0.4802 | 0.5599 | -0.6269 |
| Model 3: Input to OFC | 0.4730 | 0.5535 | -0.5518 |

A second observation concerns the OFC in conditions NE and EE. For all input mappings in these two conditions, OFC *R*^2^ values are negative which means that the models perform worse than the mean-only reference. As shown on Table 3, subjects recorded in conditions NE and EE include fewer significant voxels in their OFC compared with the NN condition. This can reduce signal quality and depress the apparent fit. This pattern could show lower task engagement of the OFC under emotional conditions or it could indicate that the voxel selection was conservative and excluded informative OFC voxels. It is important to note that fewer voxels does not necessarily imply a minor role in the network. Coupling strength analysis in Section 3.3 show that they can still exert meaningful influences. However, future work should revisit the definition and thresholding of VOI for this region.

Finally, the correlation between BMS and *R*^2^ measures varies by condition. In NN and NE, they agree for, that is the BMS-selected model also achieves the highest *R*^2^. However, in EE, *R*^2^ values are similar across model structures for each node. These results point to greater ambiguity and variability in the EE condition which can point to increased interference or noise.

### 3.3 Model validation: System properties

In this section, we use analytical tools in Section 2.5 in order to assess the system properties of the selected models. These investigations allow us to rely on the system properties for further analysis in the next section. It is worth mentioning that the model we identified is composed of a deterministic part with an additive stochastic component. In our analysis, we focus on the deterministic counterpart of this model which stands as the mean dynamics, and its stability is necessary for its generalization to the stochastic model.

#### Stability: Verification of the unforced model

It would not be biologically plausible for neuronal activity to exponentially diverge to infinity. This means that dynamical models of brain activity should be stable. Consider deterministic counterpart of the identified model in (5), i.e.,

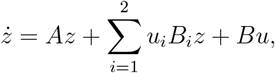

where *u*_*i*_ ∈ *{u*_1_, *u*_2_*}* such that *u*_1_ corresponds to the input of the first association of each of the conditions NN, NE, or EE, and *u*_2_ to the second one. We verify the stability of the unforced, i.e., *u*_*i*_ = 0, and forced network dynamics. In what follows, we show that the unforced system corresponding to each of the conditions, NN, NE, and EE, are asymptotically stable, see Section 2.5.

##### Property 1

*The system matrix A corresponding to the model of each of the conditions NN, NE, and EE is asymptotically stable*.

The above property verifies the validity of the identified unforced systems. We note that the values of elements of *A* matrices across the three conditions are close to each other. In order to provide a visible relation, below an average matrix is computed and sketched. The average couplings are calculated as,

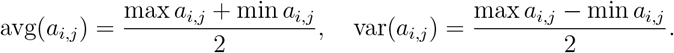

These coupling weights are illustrated on Figure 5. In this diagram, the line thickness defines the relative connection strength and the line colors indicate the sign with green as positive and red as negative. It is noteworthy that each of the directional couplings between two nodes has a comparable magnitude, representing an undirected graph. However, the pairs differ notably from one another, with Amy-Hip coupling being the strongest connection, and Hip-OFC the weakest. Finally, the largest couplings are associated with self-loops, particularly for the amygdala, indicating strong self-inhibition.

**Figure 5.**
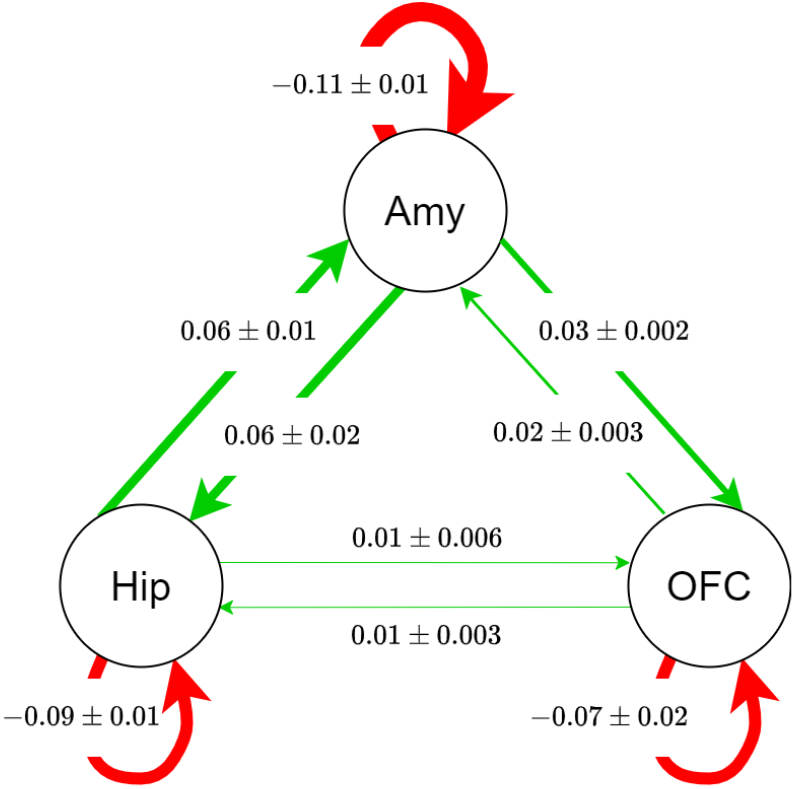
Values of elements of matrix *A* across the conditions NN, NE, EE.

#### Stability and controllability: Verification of the bilinear model

Let us now consider the forced bilinear model. We now verify stability and controllability of the identified model. Verification of these properties allows us to infer that the identified model satisfies the expected dynamical properties, hence, it can be trusted in further analysis, for instance, perturbation analysis.

Let us first verify controllability. A controllable system allows deriving its states to the desired values. Basically, it allows the level of neural activities to be controlled, for instance by regulating the coupling strengths. We perform a controllability test by checking the rank of the controllability matrix corresponding to the model of each of the three conditions. The rank of all the corresponding matrices is equal to the system dimension, three. Thus, all the models are controllable.

The dynamics of the forced model requires stability analysis with respect to its input. We verify the Input-State Stability (ISS) of the system as mentioned in Section 2.5. We first do the analysis for the overall system considering |*u*_1_| ≤ 1, |*u*_2_| ≤ 1. Note that based on the experiment, the inputs *u*_1_ and *u*_2_ are not simultaneously present. Therefore, the model in (5) can be described as 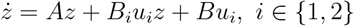.

##### Property 2

*The bilinear model* 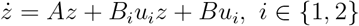 *is controllable. It is also input-to-state stable for all u*_*i*_ : |*u*_*i*_| ≤ 0.7.

Although the original unforced dynamics are asymptotically stable, input levels affect the stability when the system experiences external stimulation. We note that in the simulation results, *u*_*i*_ = 1 is used in the stochastic model which is switching between unforced and forced modes with two inputs. Above, we have characterized the condition under which the deterministic counterpart of the identified model in all modes is ISS. This condition is more conservative than the simulation where the inputs are applied for a limited time, the model is subject to noise, hence, a loss of asymptotic boundedness does not have a visible impact in simulation. For the following connectivity analysis, we set *u*_*i*_ = 0.7 for mathematical correctness.

### 3.4 Analysis: Network connectivity and coordination

After model validation, we perform connectivity analysis on the verified model to reveal the mechanisms of emotional information encoding. Consider the dynamics of the network in the presence of one of the visual stimuli, we have,

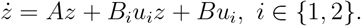

Let us denote the system equilibrium, i.e., the value of *x* for which 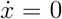 holds, by *x*^*^. Define the error variable *e* = *x* ^*−*^ *x*^*^. We obtain,

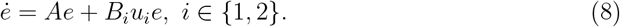

As the inputs directly correspond to the stimulation by images, the model can be thought of a switched system that switches between an unforced and a forced system, with two modes *u*_1_ or *u*_2_, every 2.5*s*. Merging the intrinsic dynamics represented by matrix *A* with the effect of the extrinsic input, we can define the system matrix as

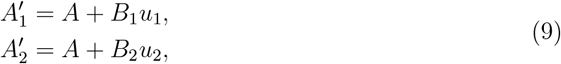

Where 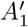 captures the dynamics when image ‘B’ is shown (*u*_1_ = 1, *u*_2_ = 0) for 2.5*s*, and 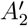 for image ‘C’ (*u*_1_ = 0, *u*_2_ = 1). The elements of *A*^*′*^ matrices are provided in Table 7 in Appendix.

Now, we have two linear systems, each corresponding to one of the inputs, and our purpose if to analyze the network connectivity and synchrony in the neuronal activities corresponding to each node. To this aim, we benefit from graph theory tools as reviewed in Section 2.5. Let us define the graph Laplacian matrix as follows

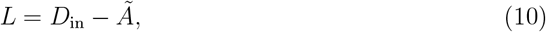

Where 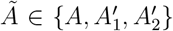 is the adjacency matrix of the network and *D*_*in*_ is the diagonal node in-degree matrix. As known, the Laplacian algebraic connectivity *λ*_2_ characterizes the network’s coordination: larger *λ*_2_ implies stronger coupling and faster consensus between neuronal activities. The network dynamics switches between input modes, and as shown in [34], the convergence rate is bounded by the *λ*_2_ values of the constituent graphs considering the switching schedule. We therefore examine *λ*_2_ for *A* (baseline), *A*_1_ (first cue) and *A*_2_ (second cue) as reported in Table 5.

**Table 5.** Algebraic connectivity *λ*_2_ in graphs and subgraphs for all conditions when no input is presented (adjacency matrix A), when *u*_1_ is active (adjacency matrix 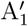), and when *u*_2_ is active (adjacency matrix 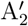). The arrows show the dominant direction of connectivity between each two pair in each condition.

| $\lambda_2$ of Graphs/Subgraphs | | | | |
| --- | --- | --- | --- | --- |
| Condition NN |  |  |  |  |
|  | Amy→Hip | Amy←OFC | Hip←OFC | Entire Graph |
| A | 0.1046 | 0.0461 | 0.0128 | 0.0411 |
| $A'_1$ | 0.1077 | 0.0504 | 0.0123 | 0.02 |
| $A'_2$ | 0.1013 | 0.0214 | 0.0107 | 0.0227 |
| Condition NE |  |  |  |  |
|  | Amy→Hip | Amy→OFC | Hip→OFC | Entire Graph |
| A | 0.1457 | 0.0512 | 0.0187 | 0.0552 |
| $A'_1$ | 0.1629 | 0.0666 | 0.0381 | 0.0832 |
| $A'_2$ | 0.1456 | 0.0389 | 0.0189 | 0.0501 |
| Condition EE |  |  |  |  |
|  | Amy←Hip | Amy←OFC | Hip←OFC | Entire Graph |
| A | 0.1023 | 0.0449 | 0.0406 | 0.0620 |
| $A'_1$ | 0.0930 | 0.0293 | 0.0284 | 0.0406 |
| $A'_2$ | 0.1430 | 0.0638 | 0.0845 | 0.0976 |

The connectivity values (*λ*_2_) of the whole graph indicate that connectivity in the NN condition remains unchanged with alternating inputs. In the NE and EE conditions, connectivity differs between emotional and neutral image presentations. Relative to the unforced (baseline) network, NE shows an initial increase followed by a decrease in connectivity strength, whereas EE shows an initial decrease followed by an increase. Averaged across *u*_1_ and *u*_2_, both NE and EE exhibit higher connectivity than baseline, with the increase being larger and more robust for NE.

In addition to the whole network analysis, we also study the sub-networks. Across all conditions, the *λ*_2_ values of the Amy-Hip sub-network are highest, indicating the strongest state coordination, e.g., consensus and synchronization, with the highest speed compared with Hip-OFC and Amy-OFC sub-networks. The highest values for Amy-Hip connectivity correspond to the NE condition, while in the EE there is increased connectivity compared to the NN condition. This indicates that the introduction of emotions leads to stronger interactions between the amygdala and the hippocampus. In contrast, hip-OFC coupling possesses the lowest *λ*_2_ values, indicating weaker and less robust coordination, except for condition EE. We shall note that the accuracy of the simulated model output for OFC versus its corresponding data in condition EE is subject to error indicated by a negative *R*^2^ value in Table 4. The latter may influence the inferred couplings between OFC and the two other regions in the condition EE. Nonetheless, our analysis stays coherent across the other conditions allowing hypothesizing the weaker connectivity of Hip-OFC. Figure 6 shows the dominant direction of influence per condition, based on Tabel 7, together with the intensity of the flow, i.e., connectivity, across the three conditions. Other couplings are omitted for the sake of clarity.

**Figure 6.**
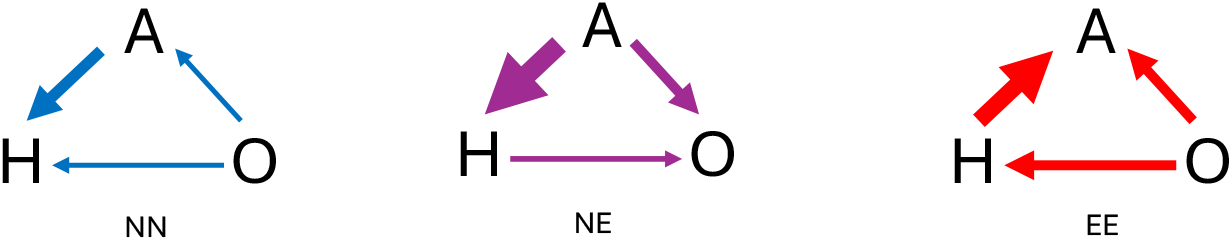
Dominant direction of connectivity in network corresponding to three conditions NN, NE, and EE.

#### Sensitivity of the model and prediction

We have analyzed network connectivity per condition using graph-theory measure, i.e., the algebraic connectivity, and showed that introducing emotional arousal increased overall network connectivity, particularly hippocampus–amygdala coupling. This effect is observed strongest in the NE condition. An interesting question is to determine the effects of changing the intensity of the emotional cues. We verify this question by varying the input signals and test system properties and network connectivity. In other words, we verify changes in system’s stability and the network’s algebraic connectivity by increasing the values of *u*_1_ and *u*_2_. We now provide a group-wise analysis by assuming perturbations of the input signals.

Condition NN: the upper bound of *u*_1_ for guaranteeing the stability of 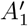 is even smaller than the threshold for other inputs. For *u*_2_, the input can be increased two times to keep the network stability, however, matrix 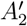 has mixed positive and negative diagonal and off-diagonal elements which does not assure an improved network connectivity.

Condition NE: considering matrix *B*_1_ corresponding to *u*_1_, we can increase the input by 10 times before 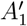 loses stability. However, the structure of the corresponding Laplacian matrix changes for *u*_1_ ≥ 6.5. Thus, the network connectivity increases up to *u*_1_ = 6.5. Considering *u*_2_, a slight increase in the value of *u*_2_ leads to instability of 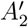.

Condition EE: When *u*_2_ is increased, 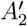 becomes unstable. In contrast, increasing *u*_1_ by fourfold keeps 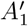 stable, although measured connectivity declines as input increases.

Our model then predicts that in the NE condition, making the first image more neutral, as a manner of increasing the intensity of *u*_1_, increases the coupling strength, hence, we infer a more enhanced memory. Also, in the EE condition, if the first emotional trigger is less intense compared with the second one, the encoding is enhanced. Looking at both of the conditions, the result is consistent: the more the first trigger carries less emotional valance compared with the second one, the greater level of integrated activities in Hip-Amy occur leading to improved associated memory.

## 4 Discussion

Here, we discuss the main objectives, modeling and mechanistic understanding by analysis, of the article by shedding a light on the obtained results and discuss limitations and future perspectives.

We have explored several dynamic models and found that a stochastic bilinear state-space model is the best candidate for emotional memory encoding in all three conditions, Neutral-Neutral, Neutral-Emotional and Emotional-Emotional. We have used control theory tools, such as input-state stability, and controllability, to validate the deterministic counterpart of the model. We then measured graph algebraic connectivity of the networks which is an indicator of the network connectivity and synchrony of neuronal activities among the amygdala, hippocampus, and OFC. Our results confirm the prior findings in [39] that memory was enhanced in the Neutral-Emotional condition, as we showed the increase of the hippocampus–amygdala coupling and overall network connectivity in this condition. Furthermore, analyzing the input-state stability of the model of Emotional–Emotional condition, we discussed that the coupling of hippocampus and amygdala, as well as the whole network connectivity increase when the first image’s valence is substantially less negative rated than the second image, but decrease otherwise. This pattern mirrors the Neutral–Emotional condition, where the first image is emotionally neutral compared with the second. Moreover, our effective connectivity analysis suggested that the amygdala predominantly influences the other two regions in the Neutral–Emotional condition, whereas the OFC plays a dominant role in the other two conditions. In what follows we further elaborate on the model, analysis, limitations and perspectives.

### Stochastic bilinear network model

As discussed in the modeling part, stochastic bilinear model served as the best fit to the experimental data of emotional encoding. In fact, stochastic DCM includes nonlinearities via the input-dependent bilinear term, and in addition, it incorporates the unmodeled and noisy elements via the stochastic component. One of the most crucial differences between the stochastic DCM and the other DCM variations examined in this work is the use of generalized coordinates which allows tracking several time derivatives of each estimated parameter, as opposed to considering only point estimates. Consequently, much more information about the shape of the function is considered, allowing the model to correctly represent vastly more complex signals.

Moreover, experimental evidence supports including stochastic uncertainties in our modeling. Neural noise tends to increase when the environment diverges from a person’s predictions [12]. In our task, images were shown rapidly and evoked highly subjective emotional responses; each valence was presented for about 8 seconds, a duration comparable to the hemodynamic response time. We note that fMRI estimates neural activity indirectly by measuring the BOLD signals which comparably posses slower dynamics. Therefore, increasing deterministic nonlinear components to capture changing dynamics is not appropriate, as reflected in our modeling choices. Stochastic DCMs accommodate these temporal fluctuations by linking fast neuronal dynamics to the slower BOLD signal.

### Analysis for revealing mechanisms

We used graph theory and stability theory tools in order to analyze the modeled dynamics and reveal its mechanisms.

Our analysis on network coordination show dependency on input context, that is, the order of the emotional valence of the images, i.e., neutral or emotional, and their intensity shape the associative memory encoding dynamics. We showed that across conditions the Amy-Hip pair consistently exhibits higher algebraic connectivity than other pairs. This result supports the view that the limbic amygdala–hippocampus loop anchors emotional associative encoding. We observed that the algebraic connectivity of the whole graph is higher in conditions containing emotional cues, i.e., Neutral-Emotional and Emotional-Emotional, consistent with emotion-induced encoding. This increase is not persistent and differs in nature between these two conditions. In Neutral-Emotional, a more intense neutrality before an emotional cue increases network connectivity and coordination, which we interpret as greater enhanced memory. In Emotional-Emotional, a milder first emotional valence is associated with higher network connectivity and integration; increasing the first image’s emotional valence appears to reduce later recall. Both findings align and together suggest that the second emotional information should be more intense to improve information integration via Amy-Hip coupling. In addition, our modeling approach infers effective connectivity, i.e., the neuronal activity of which of the three brain regions has a greater influence on the two others. Our results indicate that, in the Neutral-Emotional condition, the amygdala is the major influencer in the network.

We showed that a combined connectivity and stability analysis allow us to determine how to tune the level of emotional valence of the cues to improve or disrupt information encoding.

### Limitations and perspectives

As a known limitation of DCM, exploring a large model space is computationally demanding. Nevertheless, our results delineate a model space that can guide future data-driven modeling of emotion-associated memory across frameworks.

We focused on group-level models. However, how well these models generalize to unseen data was not assessed because of our limited neuroimaging sample. Comparing group and individual-level models and quantifying their correspondence is an important question that should be addressed in future work. Additionally, low numbers of informative voxels in the OFC reduced predictive accuracy, suggesting that more detailed investigation of the OFC’s role in the network—potentially with larger datasets—could add to the obtained knowledge.

Another key future step is to integrate the recall phase of the experiment into the connectivity analysis to verify how connectivity during encoding relates to subsequent recall or forgetting. Finally, assessing the applicability of these results for designing regulatory mechanisms, such as precision-medicine approaches or stimulation techniques, and examining implications for learning and education are important directions for future research.

## 5 Conclusion

We studied the network dynamics composed of the hippocampus, amygdala, and orbitofrontal cortex in an fMRI emotional associative-memory task. Participants viewed image pairs that were Neutral–Neutral, Neutral–Emotional, or Emotional–Emotional. Using the dynamic causal modeling framework, we showed that a stochastic bilinear state-space model best describes the neuronal dynamics in each condition. Furthermore, we analyzed the network dynamics of each condition using tools from graph theory and stability theory and discussed the differences in strength and direction of couplings, connectivity and stability of each of these networks. Our results confirmed the prior findings that introducing emotional stimuli increases hippocampus–amygdala coupling and enhances network connectivity in the Neutral–Emotional condition. In addition, we showed that in the Emotional–Emotional condition, connectivity increased when the first image’s valence was substantially less negative than the second’s. Our analyses revealed the effective connectivity and suggested that the neuronal activity of amygdala may influence the other two regions in the Neutral–Emotional condition, whereas the OFC plays a dominant role in the other two conditions. We provided an extensive discussions about our findings and suggested future directions. Particularly, we pointed out to the utility of control theory and graph theory tools in enhancing the mechanistic understanding of cortical dynamics and their potentials for designing future regulatory mechanisms.

## Acknowledgments

The authors would like to thank W. Liu, T. Kellermann, and M. Bartzioka.

## A Nonlinear and Two-state DCM

### A.1 Nonlinear DCM

On top of the assumptions of the Classical DCM, Nonlinear DCM includes an additive extra term to incorporate the effect of the activity within a node on the connectivity of the other nodes [38] resulting in the following neuronal dynamics:

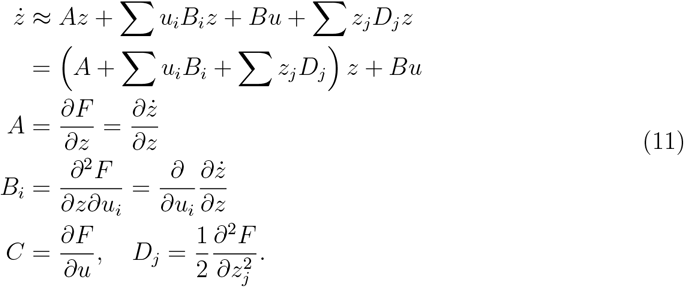

These equations are then combined with the hemodynamic state model described in Section 2.3.1 to result in the Nonlinear DCM. Within our model space of nonlinear DCM, we set the same initial values to A, B and C matrices as the classical DCM, and chose D matrices as:

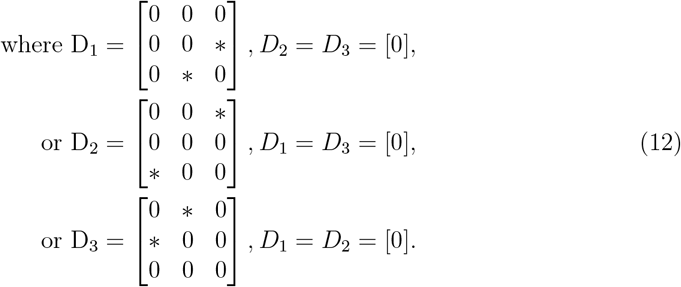

To reflect prior findings on the modulatory effect of the nodes [32, 36], we specified the *D* matrices such that only one node modulates the connectivity between the other two. Consequently, our model space comprises nine nonlinear models.

### A.2 Two-state DCM

The classical and nonlinear DCMs use a single state per node, which might not capture both the excitatory and inhibitory effects of the neuronal nodes. This limitation can be addressed with the two-state DCM which assigns 2 states per node [28], increasing the number of total states describing the neural dynamics between Amy, Hip and OFC from three to six. In this formulation, each region *i* is represented by an excitatory state 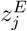 and an inhibitory state 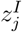. Consequently, the system’s Jacobian, previously *J* = *A* + ∑*u*_*i*_*B*_*i*_, expands to accommodate the additional state per region, resulting in:

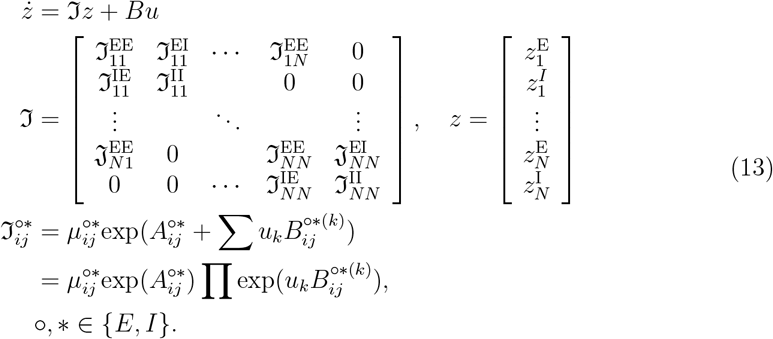

This formulation again uses the same node connections defined in Section 2.3.1 resulting in three different two-state models.

## B Estimated Parameters of the Models

**Table 6.** Estimated parameters in the optimal Stochastic DCM models for all conditions. The columns are labeled to indicate which directed connection the corresponding values belong to. “A” stands for the Amygdala, “H” for the Hippocampus, and “O” for the Orbitofrontal Cortex. All values are rounded to 4 decimal places.

| Matrix A |  |  |  |  |  |  |  |  |  |
| --- | --- | --- | --- | --- | --- | --- | --- | --- | --- |
|  | A→A | H→A | O→A | A→H | H→H | O→H | A→O | H→O | O→O |
| **NN** | -0.1033 | 0.0574 | 0.0190 | 0.0472 | -0.1093 | 0.0077 | 0.0271 | 0.0051 | -0.0416 |
| **NE** | -0.1248 | 0.0649 | 0.0191 | 0.0808 | -0.0939 | 0.0092 | 0.0321 | 0.0095 | -0.0444 |
| **EE** | -0.1015 | 0.0552 | 0.0229 | 0.0471 | -0.0782 | 0.0243 | 0.0220 | 0.0163 | -0.0902 |
| Matrix B1 |  |  |  |  |  |  |  |  |  |
|  | A→A | H→A | O→A | A→H | H→H | O→H | A→O | H→O | O→O |
| **NN** | 0.0495 | 0.0080 | -0.0060 | -0.0036 | -0.0097 | -0.0155 | 0.0122 | -0.0204 | 0.1030 |
| **NE** | -0.0757 | 0.0107 | 0.0126 | 0.0138 | 0.0003 | 0.0076 | 0.0094 | 0.0201 | -0.0407 |
| **EE** | 0.0206 | -0.0056 | -0.0121 | -0.0077 | 0.0154 | -0.0044 | -0.0101 | -0.0130 | 0.0035 |
| Matrix B2 |  |  |  |  |  |  |  |  |  |
|  | A→A | H→A | O→A | A→H | H→H | O→H | A→O | H→O | O→O |
| **NN** | 0.0002 | 0.0037 | -0.0133 | -0.0084 | -0.0336 | -0.0112 | -0.0172 | 0.0082 | 0.0246 |
| **NE** | 0.0033 | 0.0037 | -0.0140 | -0.0039 | -0.0120 | -0.0049 | -0.0036 | 0.0051 | 0.0481 |
| **EE** | -0.0411 | 0.0310 | 0.0332 | 0.0271 | -0.0201 | 0.0436 | -0.0062 | 0.0191 | 0.0130 |
| Matrix B |  |  |  |  |  |  |  |  |  |
| | $u_1 \rightarrow A$ | $u_2 \rightarrow A$ | $u_1 \rightarrow H$ | $u_2 \rightarrow H$ | $u_1 \rightarrow O$ | $u_2 \rightarrow O$ | | | |
| **NN** | -0.0042 | 0.0008 | 0 | 0 | 0 | 0 |  |  |  |
| **NE** | 0 | 0 | 0 | 0 | 0.0008 | 0.0020 |  |  |  |
| **EE** | 0 | 0 | -0.0006 | 0.0049 | 0 | 0 |  |  |  |

**Table 7.**
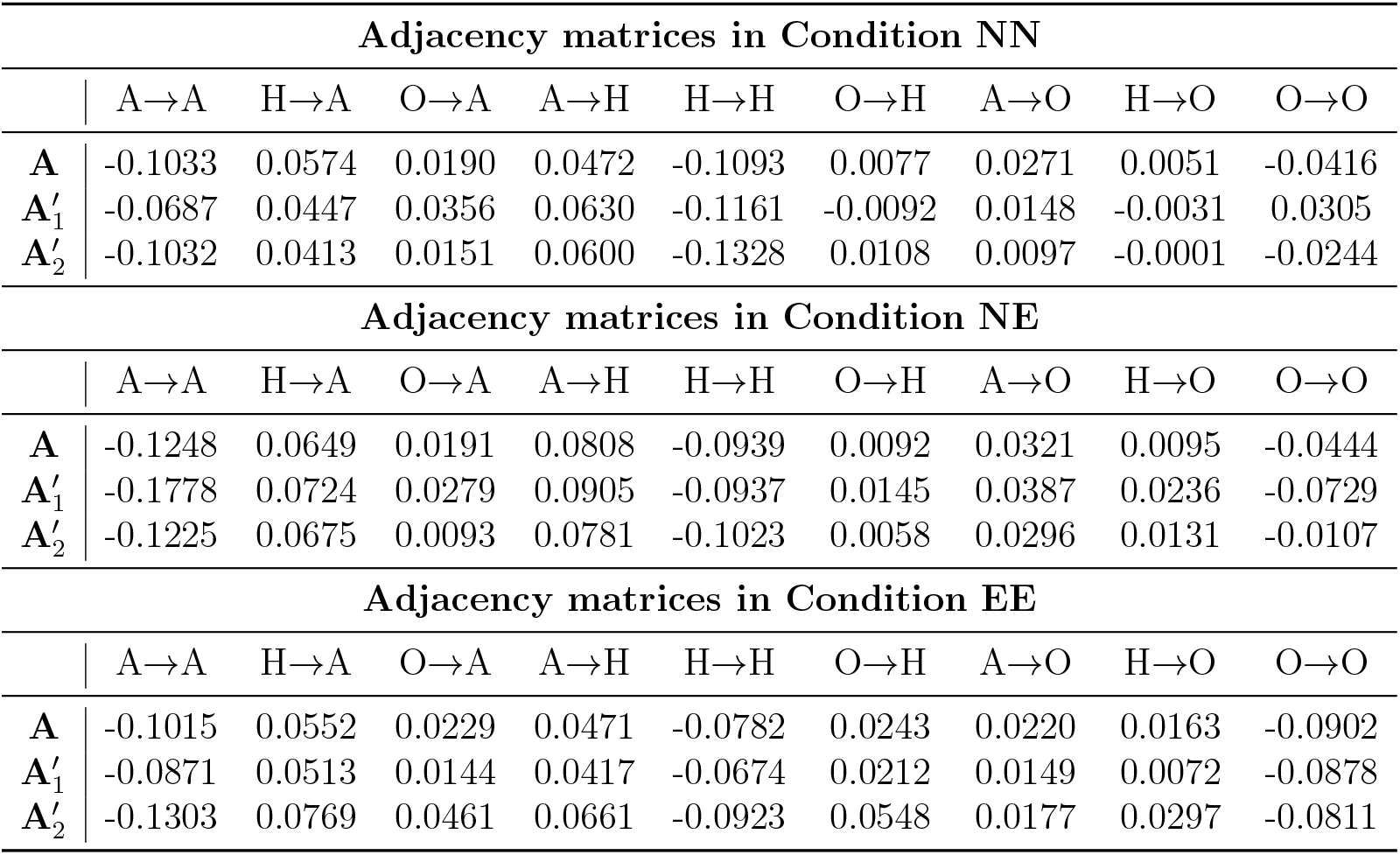
Estimated parameters of *A* and *A*^*′*^ in the optimal Stochastic DCM models for all conditions. 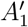 is the coupling matrix when input 1 is active, and 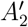 when input 2 is active. The columns are labeled to indicate which directed connection the corresponding values belong to.

## Notes

### Competing Interest Statement

The authors have declared no competing interest.

